# Novel RNA aptamers targeting gastrointestinal cancer biomarkers CEA, CA50 and CA72-4 with superior affinity and specificity

**DOI:** 10.1101/335620

**Authors:** Qing Pan, Carmen OK Law, Mingo MH Yung, KC Han, YL Pon, Terrence CK Lau

## Abstract

Gastric cancer is the third most common cause of death from cancer in the world and it remains difficult to cure in Western countries, primarily because most patients present with advanced disease. Currently, CEA, CA50 and CA72-4 are commonly used as tumor markers for gastric cancer by immunoassays. However, the drawback and conundrum of immunoassay are the unceasing problem in standardization of quality of antibodies and time/effort for the intensive production. Therefore, there is an urgent need for the development of a standardized assay to detect gastric cancer at the early stage.

Aptamers are DNA or RNA oligonucleotides with structural domain which recognize ligands such as proteins with superior affinity and specificity when compared to antibodies. In this study, SELEX (Systematic Evolution of Ligands by Exponential enrichment) technique was adopted to screen a random 30mer RNA library for aptamers targeting CEA, CA50 and CA72-4 respectively. Combined with high-throughput sequencing, we identified 6 aptamers which are specifically target for these three biomarkers of gastrointestinal cancer. Intriguingly, the predicted secondary structures of RNA aptamers from each antigen showed significant structural similarity, suggesting the structural recognition between the aptamers and the antigens. Moreover, we determined the dissociation constants of all the aptamers to their corresponding antigens by fluorescence spectroscopy, which further demonstrating high affinities between the aptamers and the antigens. In addition, immunostaining of gastric adenocarcinoma cell line AGS using CEA Aptamer probe showed positive fluorescent signal which proves the potential of the aptamer as a detection tool for gastric cancer. Furthermore, substantially decreased cell viability and growth were observed when human colorectal cell line LS-174T was transfected with each individual aptamers. Taking together, these novel RNA aptamers targeting gastrointestinal cancer biomarker CEA, CA50 and CA72-4 will aid further development and standardization of clinical diagnostic method with better sensitivity and specificity, and potentially future therapeutics development of gastric cancer.

## 1. Introduction

According to the statistic from World Health Organization 2015, Gastric cancer is the third most common cause of cancer-related death in the world [1]. Nevertheless, gastric cancer remains difficult to cure due to the lack of early detection biomarkers and most patients are diagnosed at late stage or with advanced disease. For the last decades, detection of biomarkers or tumor markers has been widely used in clinical management which assist the screening, diagnosis, prediction of prognosis and recurrence, and post-treatment monitoring. Common biomarkers for gastrointestinal cancer can be largely classified as carcinoembryonic antigen (CEA), and tumor associated antigens, such as cancer antigen 19-9 (CA19-9), cancer antigen 50 (CA50) and cancer antigen 72-4 (CA72-4) [2]. CEA, with a molecular weight of 180-200 kD, is a cell surface glycoprotein that plays a role in cell adhesion and intracellular signaling [3]. Cancer antigen 50 (CA50), with a molecular weight of ∽210 kD, is defined by the monoclonal antibody C 50 developed against a colorectal cancer (CRC) cell line COLO-205. It is elevated levels in serum and can be observed in a variety of malignancies, especially gastrointestinal cancers [4]. Cancer antigen 72-4 (CA72-4), with a molecular weight of 220-400 kD, is a mucin-like high molecular weight tumor associated antigen, and it is considered as the first choice of tumor marker for gastric carcinoma because of a superior sensitivity than CEA and CA19-9 [5].

Currently, enzyme immunoassay (EIA) and radio-immunoassay (RIA) are commonly employed for the detection of tumor markers [6]. Although the interaction between antibodies and antigens are highly specific, the biggest drawback and conundrum of immunoassays are harmonization and standardization of the assays for clinical applications [5]. Different manufactures of the commercial detection kits of tumor markers may design different binding sites/ epitopes for the immunoassays which leading to fluctuations on quality and challenge on the standardization. Moreover, variation in the induction of immune response in biological system also contribute to the uncertainty of the quality and the production is suffered from lot-to-lot variation. In addition, a minimal perturbation of conformational epitopes on native proteins will cause a failure in probing against the corresponding antigen by a monoclonal antibody. Technically, the relatively low stability and short shelf life of immunoassays also post challenge for frequent calibrations and fluctuations of the results. All these limitations lead to an urgent need for the development of a standardized method to detect tumor markers, especially gastric cancer, for clinical applications.

Aptamers which are either DNA or RNA oligonucleotides, are capable of binding different targets with high affinity and selectivity. The method for screening specific aptamers is known as ‘SELEX’ (Systematic Evolution of Ligands by Exponential enrichment), which was first developed in 1990. This ‘*in vitro* selection’ technique allows a simultaneous screening of individual nucleic acid molecules up to 10^15^ per selection for a particular target [6, 7]. Although a selection between aptamers and a target could be carried out and optimized under any condition, aptamers can be applied not only to diagnosis, but also to therapy[8, 9] such as Pegaptanib, which is an FDA approved anti-vascular endothelial growth factor (anti-VEGF) RNA aptamer for the treatment of an age-related macular degeneration [10]. The increasing interest in aptamers attributes to their versatility and superior properties. Aptamers are in general more stable than antibodies, and modification of aptamers can greatly enhance their functionality and extend their shelf life. The quality of aptamers in terms of reliability and reproducibility is more consistent because their synthesis and purification are robust under controllable settings in a machine. Furthermore, the selection and optimization of aptamers for large scale production are more simple and efficient than producing specific monoclonal antibodies. Thus, the selection for aptamers and their consequent application are rather promising in biological research and medical services.

Over the last decade, as the rapid progress of SELEX, developing aptamers for commercial and clinical interesting targets is becoming more and more efficient [11, 12]. SELEX has been successfully integrated in microfluidic chip based systems to screen aptamers against the protein targets with high affinities [11-18]. Moreover, Cell SELEX explores the expression of cell surface epitopes and distinguishes between different types of target cells [19-22]. High-throughput next-generation sequencing (NGS) and subsequent computational data analysis have also been integrated in SELEX to optimize aptamer discovery at a high resolution[11]. Sequencing reads are clustered and ranked in the order of frequencies. Candidate aptamers are further selected by a cut-off of read count [19, 23-27].

An increasing number of DNA/RNA aptamers targeting gastrointestinal cancer biomarkers and cancer cells are discovered in recent years. RNA aptamer YJ-1 binds specifically to CEA-positive cells and inhibited homotypic aggregation, migration and invasion by CEA-positive cells in mice[28]. Moreover, two DNA aptamers bound to lgV-like N domain of CEA were found inhibiting cell adhesion properties of cancer cells [29]. DNA aptamer cy-apt20 was found targeting human gastric carcinoma AGS cells while showed minimal recognition to normal gastric epithelial GES-1 cells [22]. Eight colorectal cancer stem cells (CR-CSCs)/CRC-specific aptamers were reported and three of them showed high affinities towards their respective target cells [20]. However, aptamers targeting other gastrointestinal cancer biomarkers such as CA72-4 haven’t been reported yet.

In this study, we set out to screen the library of aptamer by SELEX for three biomarkers of gastrointestinal cancer, CEA, CA50 and CA72-4, respectively. We identified 6 novel RNA aptamers (two aptamers for each biomarker) using high-throughput sequencing technology and then measured their affinity by fluorescence spectroscopy. Their dissociation constants ranged from 16.5 to 156nM which implicates high affinities between the aptamers and the antigens. Intriguingly, the predicted secondary structures of RNA aptamers from each antigen showed significant structural similarity and immunostaining using CEA aptamers demonstrated positive fluorescent signal on gastric carcinomas AGS cells, suggesting the structural recognition between the aptamers and the antigens. Moreover, we found that the cell viability and growth rate of human colorectal cell line LS-174T were substantially decreased after transfected with the aptamers which may demonstrate a functional role in cancer development. These results suggest the potential clinical applications of these aptamers as a diagnostic or therapeutic tool for gastric cancer in the future since the nature of aptamers can overcome the limitations of immunoassays with better sensitivity and specificity.

## 2. Materials and methods

### DNA oligos and antigens

An initial DNA library that had a random 30-nucleotide sequence (N30) in between as the viable region, were deliberately designed and ordered (Life Technologies) as the template (Fig 1a). The length of N30 sequence is 82 nucleotides (nt), including a T7 promoter at the 5’-end, a *Bam*H I and a *Hin*d III restriction site flanking as the boundary, and a barcode sequence at the 3’-end for discrimination in sequencing. Gastrointestinal cancer biomarkers (Human source, high purity) were respectively purchased: CEA (Abcam), CA50 (BIO-RAD), and CA72-4 (BIO-RAD).

**Fig 1.**
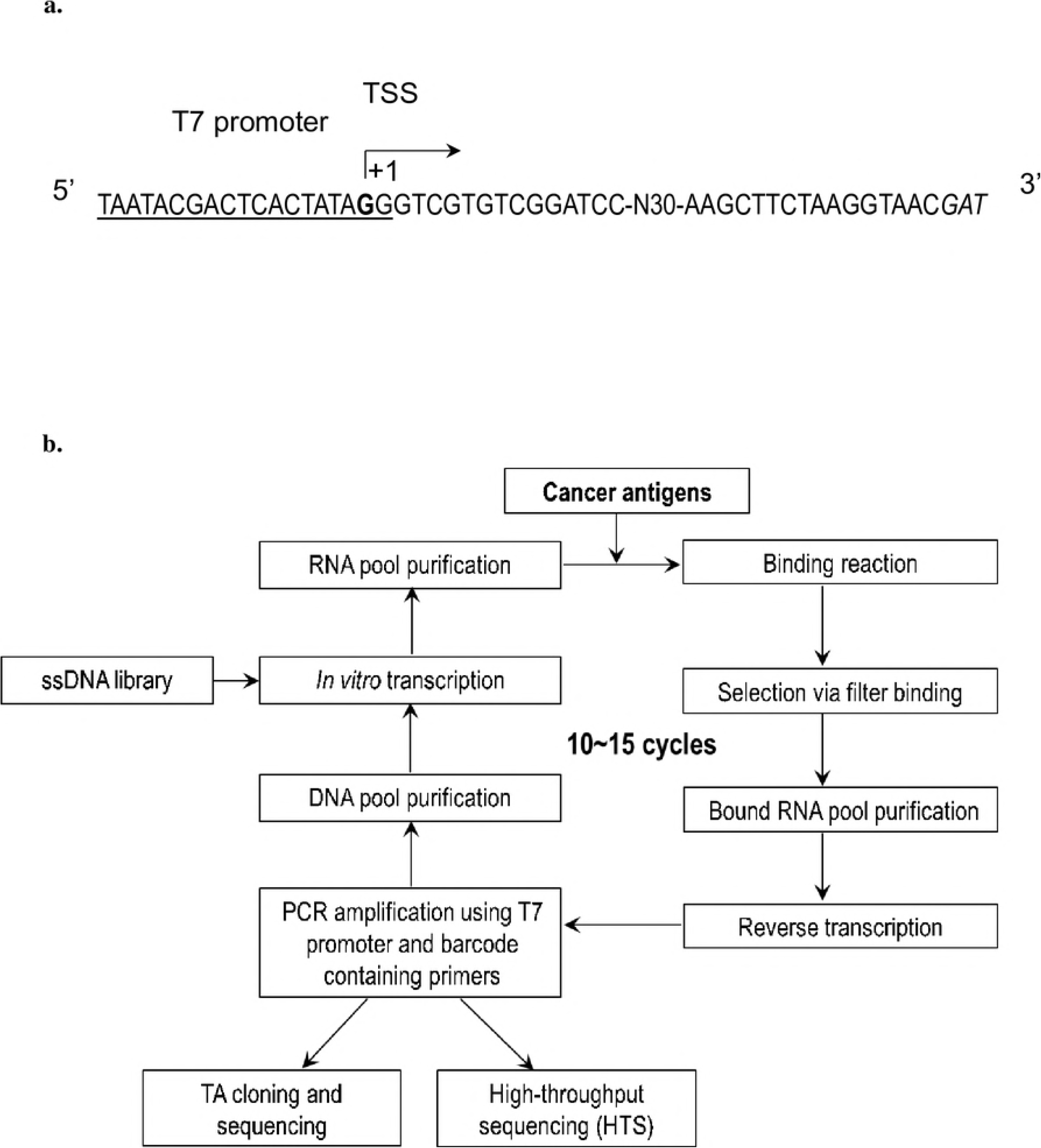
Designs of initial ssDNA library and SELEX workflow. **a)** Design and components of initial ssDNA library, underlined sequence at the 5’-end is the T7 promoter, the G in bold indicates the transcription start site (TSS), N30 refers random 30 nt in the viable region, the italic letters at the 3’-end are barcode adaptor for HTS; **b)** Workflow diagram of SELEX for screening aptamers, starting from a ssDNA library, ending at sequencing.

### SELEX and sequencing

SELEX, workflow shown in Fig 1b, was employed to screen RNA aptamers for all three gastrointestinal cancer biomarkers. The initial ssDNA library (5’-ATCGTTACCTTAGAAGCTT-N30-GGATCCGACACGACCCTATAGTGAGTCGTATTA-3’) was first transcribed into a starting RNA pool by pairing the T7 Initial-F primer (5’-AATACGACTCACTATAGGGTC-3’) and using *in vitro* transcription with T7 MEGAscript T7 Kit (Ambion) at 37°C for 6 hours. RNA product was subsequently purified by flowing through a Centri-Spin 20 column (PRINCETON). The starting RNA pool (∽10^14^ sequences) was then incubated with a cancer antigen (∽1 μg) in RSB-100 buffer (10 mM Tris-HCl, 5 mM MgCl_2_, 100 mM NaCl, 0.01% NP-40, pH 7.5) at room temperature for 30 min. Filter binding assay was performed to retain RNA-bound proteins on a PVDF membrane (GE Healthcare). Bound RNAs were released after the antigen was digested by proteinase K (Promega), and recovered through acid:phenol chloroform extraction and isopropanol precipitation. Recovered RNAs were subsequently amplified by RT-PCR using Superscript III First-Strand Synthesis System (Life Technologies) and iProof DNA polymerase (BIO-RAD) for 16-20 PCR cycles, followed by ethanol precipitation and *in vitro* transcription to transcribe dsDNA library to RNA library again to complete a SELEX cycle. Generally, the enrichment process was iteratively conducted for 10-15 cycles for selection of aptamers with elevated stringency through lowering antigen concentration, shortening incubation time, and increasing salt concentration of wash buffer. The final round of SELEX was stopped after RT-PCR. The corresponding purified DNA product was subsequently subjected to TA cloning with pGEM-T easy vector (Promega) and SANGER sequencing (BGI), as well as high-throughput sequencing (HTS) by ion Torrent sequencing platform (Life Technologies) when the RNA pool was enriched to a certain level. The quality and quantity of the same pool of DNA for ion Torrent was first verified by Agilent 2100 Bioanalyzer (Agilent Technologies). Such DNA pool was then applied to construct an amplicon library with the Ion Plus Fragment Library Kit (Ambion) according to manufacturer’s instructions, followed by sequencing on Ion 314 Chips (Life Technologies). Through a systematic bioinformatics analysis on the reads by Cygwin (Redhat), sequences were extracted and listed. Top ranking aptamers for each cancer biomarker were chosen for following biochemical characterization.

### Purification of RNA aptamers

To characterize top ranking aptamers, specific full length oligos, with replacement of N30 in the initial DNA library by identified sequences, were chemically synthesized for further study. Similarly, RNA aptamers were first generated through *in vitro* transcription using T7 MEGAscript Transcription Kit (Ambion), and purified on a 6% denaturing urea/polyacrylamide gel. The corresponding bands were visualized under UV, excised and chopped. RNA aptamers were eluted with 400 μl RNA elution buffer (0.5 M ammonium acetate, 1 mM EDTA, 0.1% SDS, pH 8.0) at 42°C overnight. Purified RNA aptamers were recovered through acid:phenol chloroform extraction, isopropanol precipitation, and dissolving in appropriate volume of RNase-free distilled water.

### Determination of binding profile

RNA aptamers were fluorescence labeled with Ulysis Alexa Fluor 488 nucleic acid labeling Kit (Life Technologies) following manufacturer’s instructions. Briefly, 1 μg ethanol precipitated gel purified RNA aptamers were mixed with 1 μl ULS reagent in DMSO and added up to 25 μl with labeling buffer. The mix was incubated at 90°C for 10 min, followed by stopping the reaction on ice. A Centri-Spin 20 column (PRINCETON) was applied to purify fluorescence labeled RNA. Next, samples for RNA-antigen interaction were prepared by incubating 1 pmol fluorescence labeled RNA and titrated with various concentrations of antigen (0, 10, 20, 50, 100, 200 nM) in RSB-100 buffer in the presence of non-specific competitor yeast tRNA (1ug/ul) at room temperature for 10 min respectively. Excitation fluorescence values at 516 nm were measured using Fluormax-4 Spectrofluorometer (Horiba), and fluorescence changes were obtained by subtracting the value in the absence of antigen [30]. Binding profiles were plotted and equilibrium dissociation constant K_*d*_ values were calculated using SigmaPlot v11.0 software (Systat).

### Cell line and cell culture

The human colon adenocarcinoma cell line LS-174T and gastric adenocarcinoma cell line AGS used in this study were purchased from American type Culture Collection (ATCC). LS-174T cells were grown and maintained in culture flask with Dulbecco’s Modified Eagle Medium (DMEM) (Life Technologies) supplemented with 10% fetal bovine serum (FBS) and 1% penicillin/streptomycin (PS), incubating at 37°C in a 5% CO_2_ environment. For passage, cells were detached by trypsin, spun down, and resuspended in fresh DMEM with FBS and PS, followed by a 1/10 subculture in flask. AGS cells were grown and maintained in RPMI-1640 medium 10% FBS and 1% penicillin/streptomycin (PS), incubating at 37°C in a 5% CO_2_ environment.

### Transfection with aptamers

Cell density of LS-174T was determined by trypan blue (Life Technologies) staining. Around 4×10^5^ cells were seeded in a 6-well plate to reach 50% confluence overnight, and transfected with 1 μg gel purified selected RNA aptamer in 10 μl Lipofectamine 2000 (Life Technologies) following the manufacturer’s instructions. Antibiotic-free DMEM for transfection was replace by 2 ml DMEM with FBS and PS after 5-hour incubation. The 48-hour post-transfected cells were detached by trypsin for following assays.

### Evaluation of cell viability and cell growth

Cell viability was determined by trypan blue exclusion. Briefly, 10 μl detached LS-174T cells were stained with 10 μl 0.4% trypan blue solution, and 10 μl mix was then loaded onto a hemocytometer. Numbers of viable cells (transparent) and dead cells (blue) were counted respectively under a light microscope at low magnification. Cell viability was calculated as the number of viable cells divided by the total number of cells within each 4×4 grid on the hemocytometer.

Cell growth rate was determined by MTT tetrazolium reduction assay. LS-174T cells treated with various RNA aptamers were seeded as 3000 cells/well in 96-well plates respectively. After 24, 48, 72 and 96-hour (as Day 0, 1, 2, 3) incubation, the plates were taken out one by one, and assayed with 10 μl MTT reagent mixture (5 mg/ml in PBS) incubating at 37°C for 4 hours. Absorbance at 570 nm was then measured by a Powerwave XS microplate reader (BioTek). Relative cell viability was calculated as A_570 nm on Day N_ divided by A_570 nm of Day 0_.

### Immunostaining of gastric adenocarcinoma cell line using RNA aptamer

AGS were grown on 10mm coverslip, harvested by washing with PBS for three times and fixed with cold methanol for 10 min. After washing three time with PBS, cells were incubated with binding buffer (BSA 1mg/ml in Dulbecco’s PBS without calcium and magnesium (pH 7.3), Mg_2_Cl 5mM, glucose 4.5g/L, Salmon Sperm DNA 0.1 mg/ml and yeast tRNA 0.1 mg/ml) for 30 min, followed by incubation of 1:100 RNA aptamer targeting CEA or 1:100 anti CEA antibody (Abcam) in binding buffer for 1 hr. For secondarty antibody incubation, cells was washed with PBS for three times and incubate with 1:1000 rabbit anti-mouse/FITC antibody (Abcam) for 30 min. The cells were then washed three time with buffer, mounted, sealed and observed by fluorescence microscopy. All images were captured under a fluorescent microscopy with 400X magnification.

## 3. Results

### RNA aptamers isolated for CEA, CA50 and CA72-4

A random library of N equal to 30 was used to screen RNA aptamers. The initial number of DNA template for *in vivo* transcription was set as approximately 10^14^, which means 10^14^ different RNA molecules in the starting RNA pool. This number of RNA copies which is equivalent to 2-5 μg covers the majority of the possible combinations of N30. In order to eliminate the non-specificity of the RNA to the PVDF membrane, a pre-clearing step was performed by immersing the membrane into the initially transcribed RNA pools for 15 min. The recovered RNA pool was then incubated with a cancer antigen to initiate SELEX.

Enrichment of RNA aptamers specific to cancer antigens was monitored by the binding ratio calculated before and after the binding experiment (amount of recovered RNA versus amount of input RNA). Through multiple rounds of SELEX, binding ratios of 18.1% for CEA at Round 14, 15.6% for CA50 at Round 12, and 24.6% for CA72-4 at Round 15 were recorded, indicating that specific RNA aptamers for cancer antigens were accumulated to a certain level, respectively.

In order to assess the sequences of enriched RNA aptamers after selection, the purified aptamers were reverse transcribed, amplified, overhangs addition at 3’ end and subcloned into the TA vector. For each antigen, 15-20 single colonies were randomly picked for sequencing. As shown in Table 1, the consensus sequences (*, **) were successfully identified in all three RNA pools, indicating the feasibility of the enrichment methodology and the possibility to subject for high-throughput sequencing.

**Table 1.**
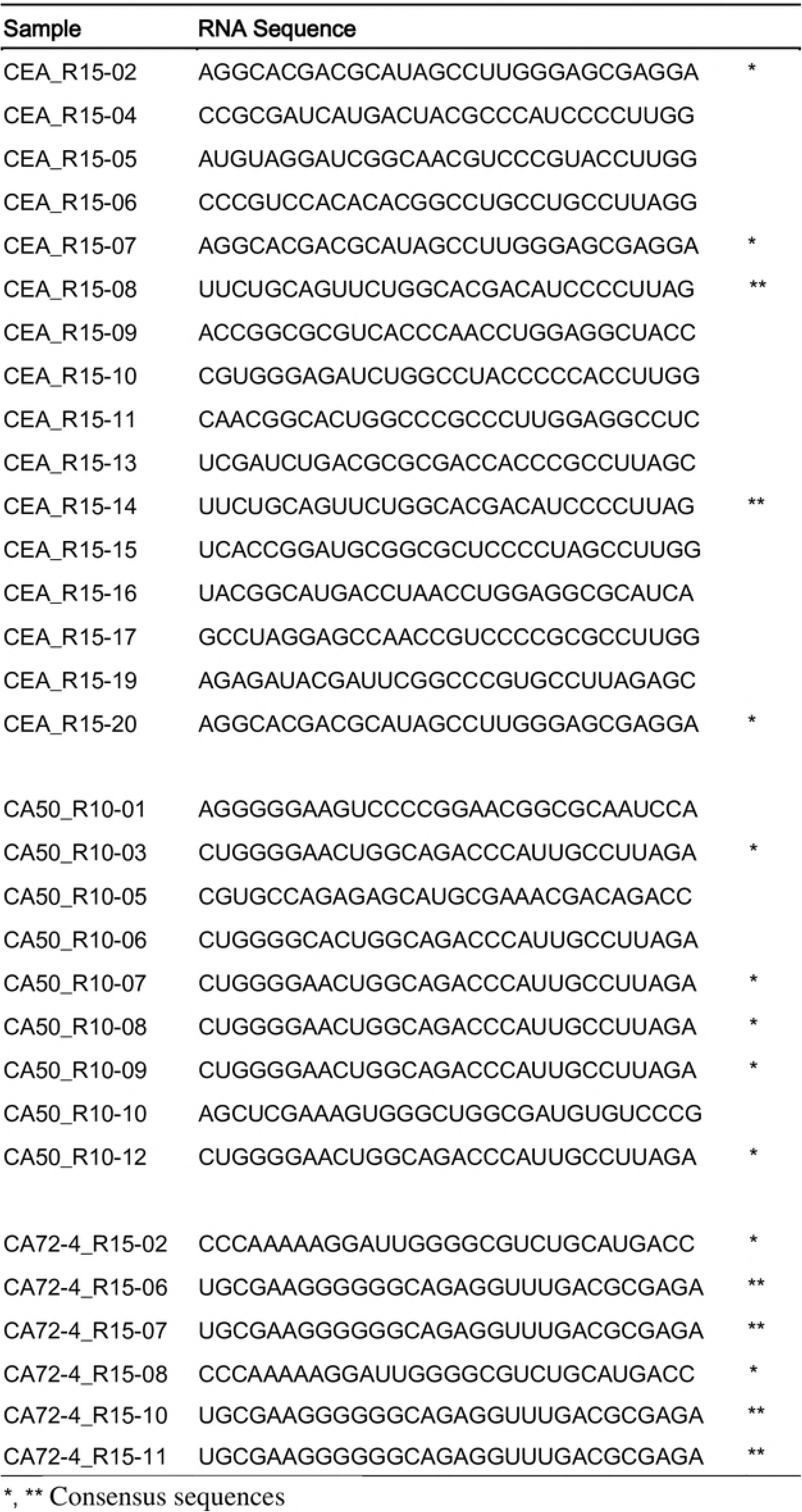
N30 RNA sequences identified by SANGER sequencing

### Highly abundant aptamers revealed by high-throughput sequencing (HTS)

Since SANGER sequencing could only provide limited data, we adopted the high-throughput sequencing (HTS) to reveal the entire population of enriched RNA aptamers for the three antigens. To do this, RNA aptamers of CEA at Round 15, CA50 at Round 12, and CA72-4 at Round 15 were subjected to HTS using ion PGM™ system. The libraries were constructed and sequenced for each antigens according to the manufacturer protocols. The sequences were sorted and ranked based on their number of reads as shown in Table 2, the distribution of the identified aptamers were shown in Fig 2. In our results, the consensus sequences revealed by SANGER sequencing were all found in top 5 ranking sequences in HTS. In order to observe the profile of the aptamers in each case, the distribution of the aptamers was plotted as a pie chart. As shown in Fig 2 and Table 3, distributions of top 5 ranking sequences of CEA, CA50 and CA72-4 were up to 40%, 80% and 90% respectively, suggesting that these consensus sequences were enriched in SELEX experiments, and became dominant in the RNA pools because of their relatively high affinity to the antigens.

**Table 2.**
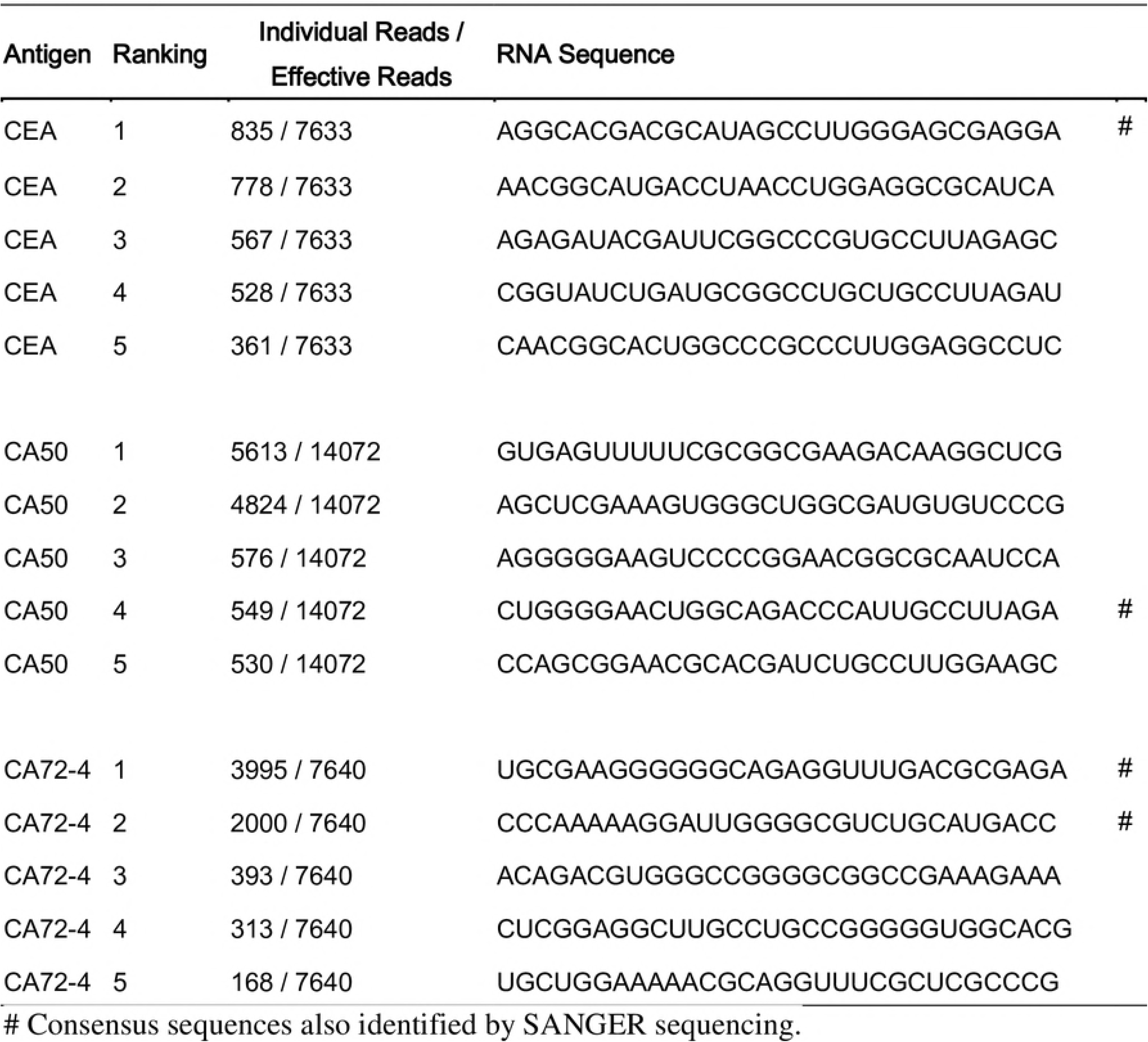
Top 5 N30 RNA sequences identified by HTS

**Table 3.**
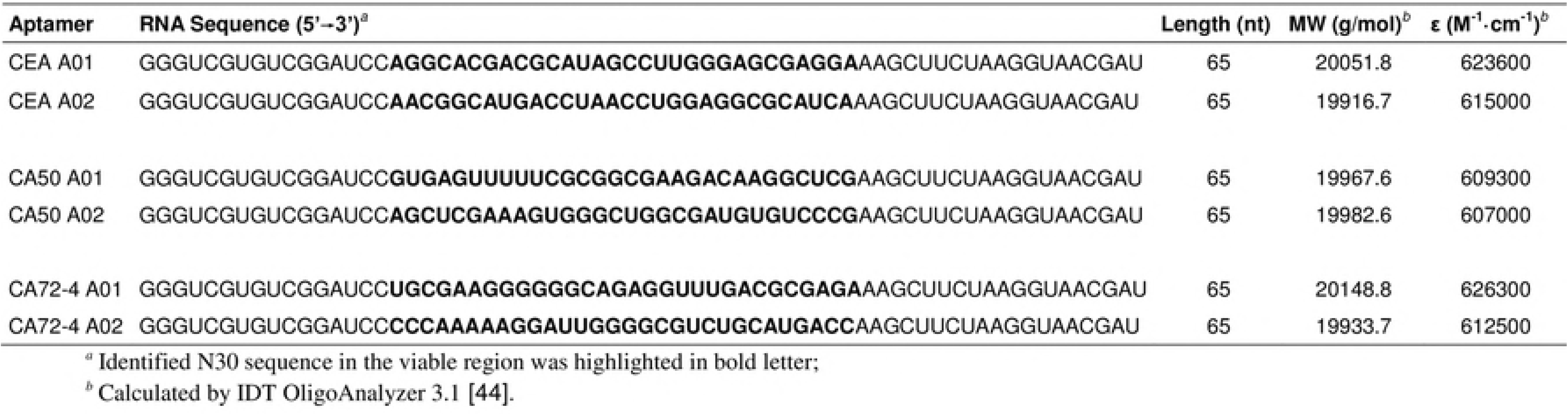
Sequences and properties of selected RNA aptamers

**Fig 2.**
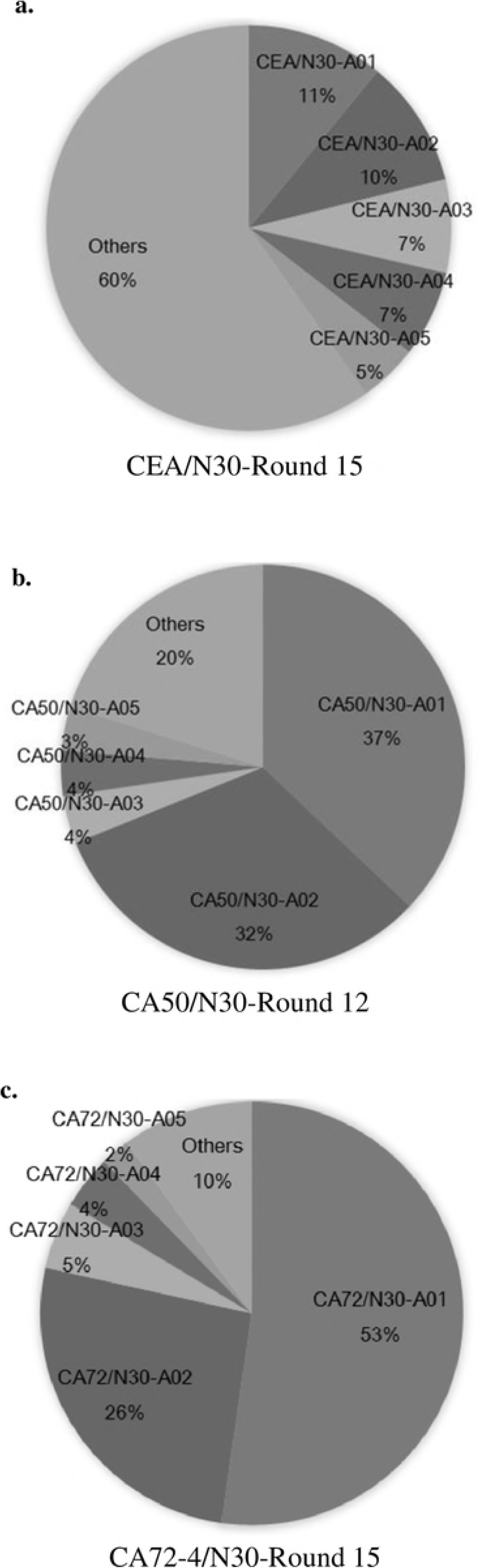
Distribution of top 5 N30 RNA sequences for CEA, CA50 and CA72-4 identified by HTS.

### Biochemical characterization of RNA aptamers

To characterize the RNA aptamers discovered from HTS, the top two RNA aptamers for each antigen were selected for further studies. The secondary structures of all six RNA aptamers, CEA A01, A02; CA50 A01, A02, and CA72 A01, A02 (Table 3), were predicted by *mfold* RNA Folding program [31]. As shown in Fig 3, interestingly, every two RNA aptamers for each biomarker exhibited similar folding and secondary structures without high sequence identity or similarity, suggesting that the aptamer structure may play an essential role in binding affinity and specificity. To evaluate these experimentally, we performed the titration assay using synthetic fluorescence labelled RNA aptamer to calculate their affinity to the corresponding antigens. The aptamers were *in vitro* transcribed, purified and labeled with fluorescence probe Alexa 488. The fluorescence labeled RNA aptamers were titrated with increasing amount of cancer antigens and the enhancement of fluorescence intensity was used to plot the binding isotherms. The curves were then fitted with the nonlinear regression-one site saturation ligand binding equation to obtain dissociation constant K_*d*_ as shown in Fig 3. Dissociation constant K_*d*_ values of CEA A01, A02 were 16.5 and 156 nM respectively; K_*d*_ values of CA50 A01, A02 was 38.0 and 30.7 nM respectively; K_*d*_ values for CA72 A01, A02 were 52.7 and 71.2 nM respectively (Table 4).

**Table 4.** Dissociation constants of the aptamer

| | Dissociation Constants, $K_d$ (nM) | Regression of the fitting |
| --- | --- | --- |
| CEA A01 | 16.5 | 0.98 |
| CEA A02 | 156.0 | 0.99 |
| CA50 A01 | 38.0 | 0.99 |
| CA50 A02 | 30.7 | 0.99 |
| CA72-4 A01 | 52.7 | 0.98 |
| CA72-4 A02 | 71.2 | 0.99 |

**Fig 3.**
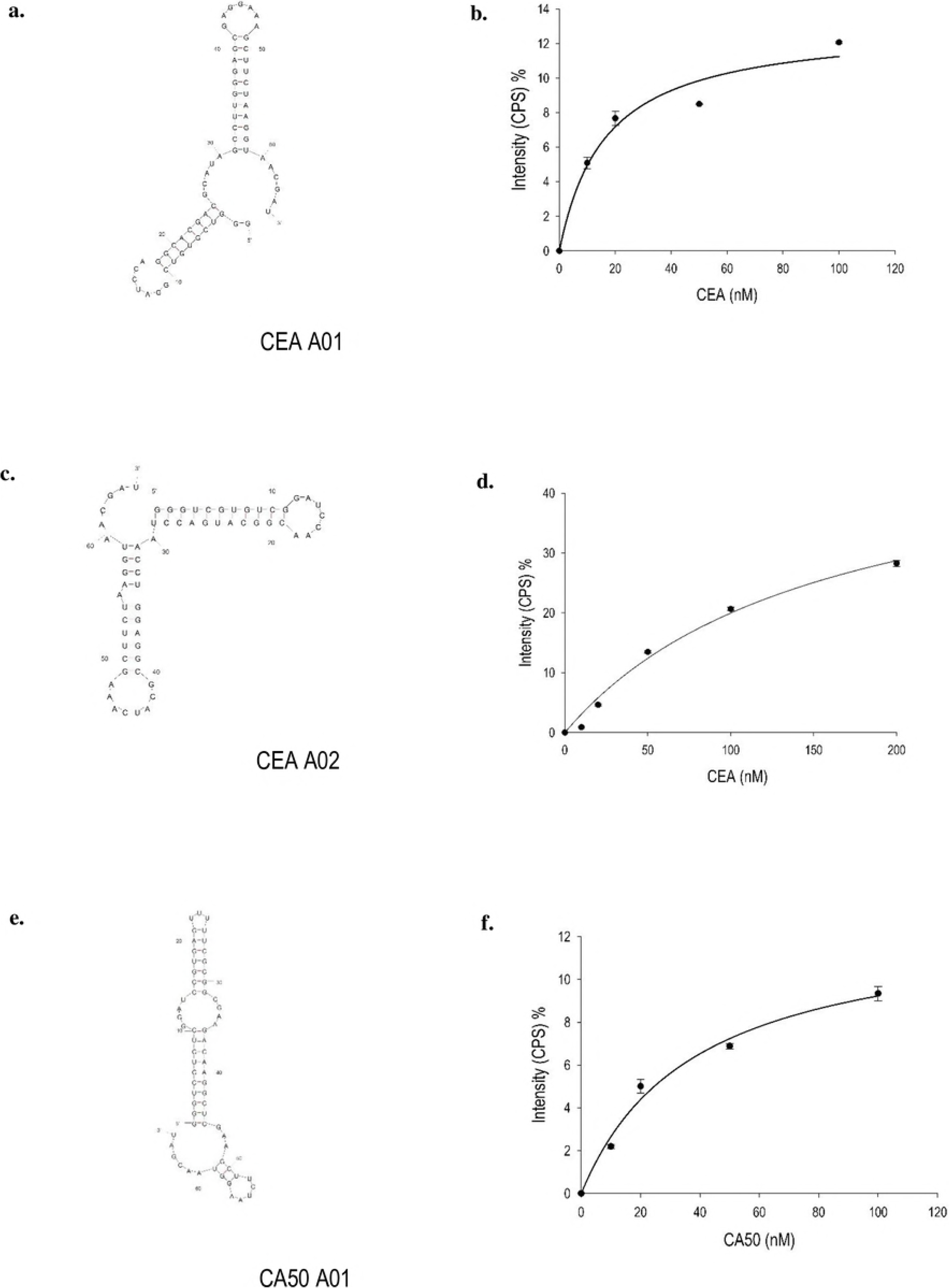
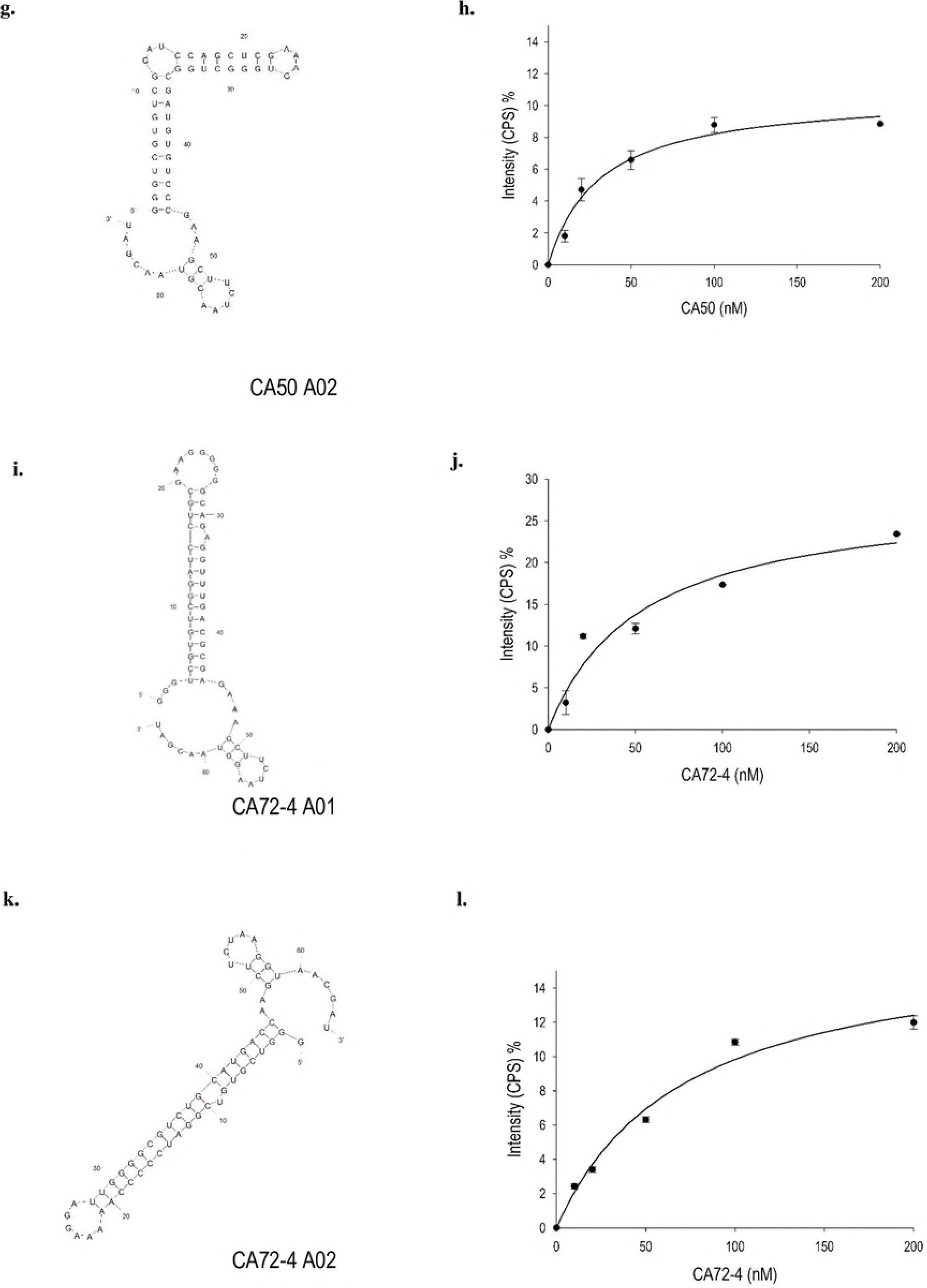
Secondary structure predicted by *mfold* [35], equilibrium dissociation curve and constant K_*d*_ of 6 RNA aptamers.

### Binding of CEA aptamers to gastric adenocarcinoma cell line AGS

To investigate if the CEA aptamers can bind to cancer cells *in vitro*, immunostaining of AGS was performed using the CEA aptamers probe. AGS cells incubated with CEA aptamers showed positive fluorescent signal same as CEA antibody (Fig 4) whereas control demonstrated no signal. The positive fluorescent signal of the immunostaining using aptamers indicated the potential application of the aptamers as the cancer detection tool and the possibility to overcome the limitation of traditional immunoassays using unstandardized antibodies.

**Fig 4.**
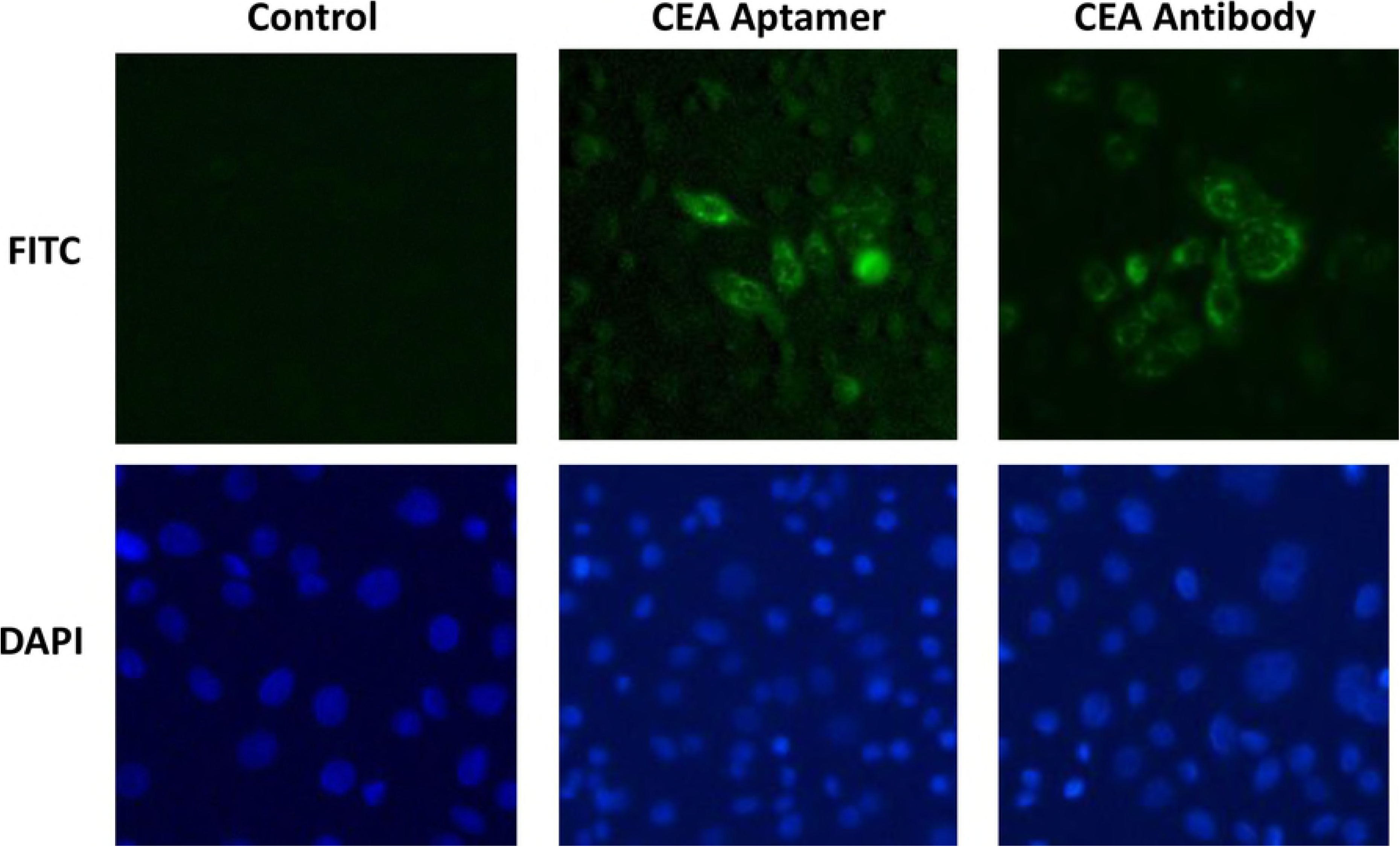
Immunostaining of AGS cells by using CEA Aptamers probe (green), CEA antibody (secondary with FITC) and DAPI (blue) stained the nucleus. All images were taken using a fluorescent microscopy with 400X magnification.

### Inhibitory effects of aptamers on LS-174T cells

To explore the potential role of RNA aptamers, we transfected LS-174T cells with selected aptamers and studied cell viability and growth rate. The most valuable virtue of LS-174T cell line is that it produced all three cancer biomarkers CEA [32], CA50 [33] and CA72-4 [34] in cell, which facilitates our study towards the comparison of all three biomarkers. Cell viability was counted 48-hour after the transfection. Viable cells of both wild-type (WT, non-transfected) and negative control (si-NC, irrelevant siRNA) retained 85% and 87% (Fig 5a), indicating the transfection alone did not affect their growth and viability. On the other hand, viability of aptamer-treated cells greatly decreased to a range from 52% to 68% of total number of cell (Fig 5a). Moreover, study on the rate of cell growth demonstrated 2-3 folds suppression of aptamer-treated cells compared with WT after 3 days (Fig 5b), suggesting that the growth of LS174T cells was also inhibited by the aptamers.

**Fig 5.**
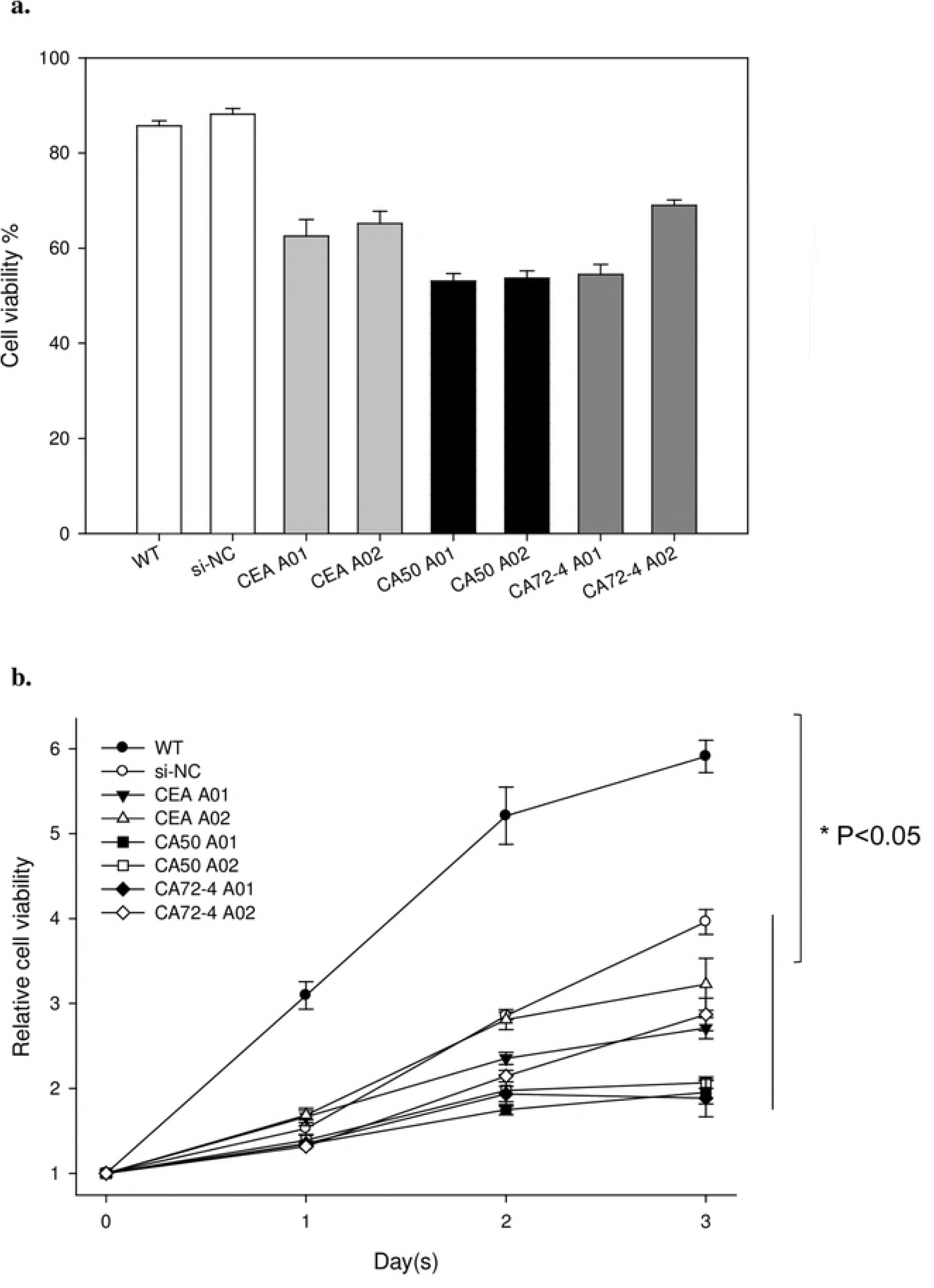
Inhibitory effects of selected aptamers on LS-174T cells. a) Cell viability determined by trypan blue staining 48-hour after the transfection; b) Relative cell growth rate monitored by the MTT assay from day 0 to day 3.

## 4. Discussion

To tackle the limitations of using immunoassays for the detection of tumor markers, we established the SELEX method incorporated with high-throughput sequencing by ion Torrent (Ion PGM™system) to screen RNA aptamers from a random RNA pool. Total 6 novel RNA aptamers for three biomarkers of gastrointestinal cancer, CEA, CA50 and CA72-4, were identified using SELEX and their biochemical properties were characterized. Although their affinities to each biomarker varied, they could reduce the viability and growth of tumor cells after transfection. Our pioneering development of SELEX to screen RNA aptamers against cancer biomarkers expanded the possibility of SELEX application as an alternative way to target such biomarkers besides antibodies.

One important improvement for our experiment is the application of high-throughput sequencing (HTS) for RNA SELEX. HTS techniques were developed and mature in recent years, which enabled researchers to screen efficiently and effectively, and profile nucleotide sequences within days. The compatibility of HTS in DNA SELEX was confirmed by several previous studies [14, 35], but it was not very commonly used in RNA SELEX. The application of HTS to inspect the RNA pool benefits us to globally view the whole population rather than an individual sequence, overcoming the shortage of insufficient information provided by limited SANGER sequencing [35]. In our study, HTS was carried out by Ion Torrent, which is a rapid, compact and economic technique for next generation sequencing. Top ranking sequences were obtained by subsequently statistical analysis, which clearly facilitate the selection process for promising aptamers. It is clear that this improvement undoubtedly opens a new pipeline to overview and designate aptamers for SELEX experiments.

The advantage of employing the human colorectal carcinoma cell line LS-174T as a model to study the potential role of RNA aptamers is that this cell line carries all three biomarkers of CEA [32], CA50 [33] and CA72-4 [34]. CEA are glycosyl phosphatidyl inositol (GPI) cell surface anchored glycoproteins, who may cooperate with CD44 variant isoforms to mediate colon carcinoma cell adhesion to E-and L-selectin [36]. CA50 and CA72 are both carbohydrate antigens with heavy glycosylations [4, 37], but their functions remained unclear. The diversity of glycosylation on cancer antigens varied their glycan compositions, molecular weights and structures [38], which might hamper the screening by SELEX for highly specific aptamers. Even so, HTS can facilitate the identification of highly abundant and most promising aptamers targeting the cancer biomarkers. The mechanism of the inhibitory effect on cell viability and growth by RNA aptamers is unclear and yet to be determined.

Taken together, this study demonstrates that SELEX incorporated with HTS is a promising and powerful tool to screen aptamers for various targets. To obtain qualified aptamers with high specificity against cancer antigens, SELEX refinement and post-SELEX development of aptamers are yet indispensable. In order to enhance the sensitivity and specificity of aptamers for future diagnostic and clinical applications, several modifications could be considered: to truncate ribonucleic acids in block to simplify the aptamer and specify the core region for RNA-protein interactions; to individually substitute / delete / insert a particular ribonucleic acid in the core region of aptamer to improve its specificity; to chemically modify particular ribonucleic acid(s) of aptamer with locked nucleic acid (LNA) to decrease its susceptibility to nucleases and increase its stability for long-term storage. Indeed, it is possible that such aptamers would become applicable in clinical applications or even therapeutics in the near future with better sensitivity and specificity, and without the limitations of immunoassays.

## Acknowledgements

This work was fund and supported by General Research Fund Early Career Scheme CityU 189813, ITF Fund 9440086, Ivorist Scientific Tech (Hong Kong) Company Limited, and Shenzhen New Industries Biomedical Engineering CO., Ltd.

## References

1. World Health Organization Cancer: Fact Sheet No 297. World Health Organization 2015, (May 21) Available at http://www.who.int/mediacentre/factsheets/fs297/en/.

2. Malati T. : Tumour markers: An overview. Indian journal of clinical biochemistry : IJCB 2007, (22) 2 17–31.

3. Terry WD, Henkart PA, Coligan JE and Todd CW. : Carcinoembryonic antigen: characterization and clinical applications. Transplantation reviews 1974, (20) 0 100–129.

4. Bunworasate U and Voravud N. : CA 50: a tumor marker for gastrointestinal malignancies. Journal of the Medical Association of Thailand = Chotmaihet thangphaet 1995, (78) 5 255– 270.

5. Sturgeon C. : Standardization of tumor markers - priorities identified through external quality assessment. Scandinavian journal of clinical and laboratory investigation.Supplementum 2016, (245) S94–9.

6. Hall B, Arshad S, Seo K, Bowman C, Corley M, Jhaveri SD and Ellington AD. : In vitro selection of RNA aptamers to a protein target by filter immobilization. Current protocols in molecular biology / edited by Frederick M. Ausubel …[et al.] 2009, (Chapter 24) Unit 24.3.

7. Kong HY and Byun J. : Nucleic Acid aptamers: new methods for selection, stabilization, and application in biomedical science. Biomolecules & therapeutics 2013, (21) 6 423–434.

8. Sun H, Zhu X, Lu PY, Rosato RR, Tan W and Zu Y. : Oligonucleotide aptamers: new tools for targeted cancer therapy. Molecular therapy.Nucleic acids 2014, (3) e182.

9. Ma H, Liu J, Ali MM, Mahmood MA, Labanieh L, Lu M, Iqbal SM, Zhang Q, Zhao W and Wan Y. : Nucleic acid aptamers in cancer research, diagnosis and therapy. Chemical Society Reviews 2015, (44) 5 1240–1256.

10. Ng EW, Shima DT, Calias P, Cunningham ET,Jr, Guyer DR and Adamis AP. : Pegaptanib, a targeted anti-VEGF aptamer for ocular vascular disease. Nature reviews.Drug discovery 2006, (5) 2 123–132.

11. Blind M and Blank M. : Aptamer selection technology and recent advances. Molecular Therapy—Nucleic Acids 2015, (4) (1) e223.

12. Ulrich H and Wrenger C. : Disease-specific biomarker discovery by aptamers. Cytometry.Part A : the journal of the International Society for Analytical Cytology 2009, (75) 9 727–733.

13. Oh SS, Ahmad KM, Cho M, Kim S, Xiao Y and Soh HT. : Improving aptamer selection efficiency through volume dilution, magnetic concentration, and continuous washing in microfluidic channels. Analytical Chemistry 2011, (83) 17 6883–6889.

14. Cho M, Xiao Y, Nie J, Stewart R, Csordas AT, Oh SS, Thomson JA and Soh HT. : Quantitative selection of DNA aptamers through microfluidic selection and high-throughput sequencing. Proceedings of the National Academy of Sciences of the United States of America 2010, (107) 35 15373–15378.

15. Oh SS, Qian J, Lou X, Zhang Y, Xiao Y and Soh HT. : Generation of highly specific aptamers via micromagnetic selection. Analytical Chemistry 2009, (81) 13 5490–5495.

16. Cho M, Soo Oh S, Nie J, Stewart R, Eisenstein M, Chambers J, Marth JD, Walker F, Thomson JA and Soh HT. : Quantitative selection and parallel characterization of aptamers. Proceedings of the National Academy of Sciences of the United States of America 2013, (110) 46 18460–18465.

17. Huang CJ, Lin HI, Shiesh SC and Lee GB. : Integrated microfluidic system for rapid screening of CRP aptamers utilizing systematic evolution of ligands by exponential enrichment (SELEX). Biosensors & bioelectronics 2010, (25) 7 1761–1766.

18. Lai HC, Wang CH, Liou TM and Lee GB. : Influenza A virus-specific aptamers screened by using an integrated microfluidic system. Lab on a chip 2014, (14) 12 2002–2013.

19. Thiel WH, Bair T, Peek AS, Liu X, Dassie J, Stockdale KR, Behlke MA, Miller FJ,Jr and Giangrande PH. : Rapid identification of cell-specific, internalizing RNA aptamers with bioinformatics analyses of a cell-based aptamer selection. PloS one 2012, (7) 9 e43836.

20. Hung LY, Wang CH, Che YJ, Fu CY, Chang HY, Wang K and Lee GB. : Screening of aptamers specific to colorectal cancer cells and stem cells by utilizing On-chip Cell-SELEX. Scientific reports 2015, (5) 10326.

21. Li WM, Bing T, Wei JY, Chen ZZ, Shangguan DH and Fang J. : Cell-SELEX-based selection of aptamers that recognize distinct targets on metastatic colorectal cancer cells. Biomaterials 2014, (35) 25 6998–7007.

22. Cao HY, Yuan AH, Shi XS, Chen W and Miao Y. : Evolution of a gastric carcinoma cell-specific DNA aptamer by live cell-SELEX. Oncology reports 2014, (32) 5 2054–2060.

23. ’t Hoen PA, Jirka SM, Ten Broeke BR, Schultes EA, Aguilera B, Pang KH, Heemskerk H, Aartsma-Rus A, van Ommen GJ and den Dunnen JT. : Phage display screening without repetitious selection rounds. Analytical Biochemistry 2012, (421) 2 622–631.

24. Schutze T, Wilhelm B, Greiner N, Braun H, Peter F, Morl M, Erdmann VA, Lehrach H, et al. : Probing the SELEX process with next-generation sequencing. PloS one 2011, (6) 12 e29604.

25. Thiel KW, Hernandez LI, Dassie JP, Thiel WH, Liu X, Stockdale KR, Rothman AM, Hernandez FJ, McNamara JO, 2nd and Giangrande PH. : Delivery of chemo-sensitizing siRNAs to HER2+- breast cancer cells using RNA aptamers. Nucleic acids research 2012, (40) 13 6319–6337.

26. Berezhnoy A, Stewart CA, Mcnamara JO, 2nd, Thiel W, Giangrande P, Trinchieri G and Gilboa E. : Isolation and optimization of murine IL-10 receptor blocking oligonucleotide aptamers using high-throughput sequencing. Molecular therapy : the journal of the American Society of Gene Therapy 2012, (20) 6 1242–1250.

27. Huang YZ, Hernandez FJ, Gu B, Stockdale KR, Nanapaneni K, Scheetz TE, Behlke MA, Peek AS, et al. : RNA aptamer-based functional ligands of the neurotrophin receptor, TrkB. Molecular pharmacology 2012, (82) 4 623–635.

28. Lee YJ, Han SR, Kim NY, Lee SH, Jeong JS and Lee SW. : An RNA aptamer that binds carcinoembryonic antigen inhibits hepatic metastasis of colon cancer cells in mice. Gastroenterology 2012, (143) 1 155–65.e8.

29. Orava EW, Abdul-Wahid A, Huang EH, Mallick AI and Gariepy J. : Blocking the attachment of cancer cells in vivo with DNA aptamers displaying anti-adhesive properties against the carcinoembryonic antigen. Molecular oncology 2013, (7) 4 799–811.

30. Jiang Y, Fang X and Bai C. : Signaling aptamer/protein binding by a molecular light switch complex. Analytical Chemistry 2004, (76) 17 5230–5235.

31. Zuker M. : Mfold web server for nucleic acid folding and hybridization prediction. Nucleic acids research 2003, (31) 13 3406–3415.

32. Shi ZR, Tsao D and Kim YS. : Subcellular distribution, synthesis, and release of carcinoembryonic antigen in cultured human colon adenocarcinoma cell lines. Cancer research 1983, (43) 9 4045–4049.

33. Frykholm G, Glimelius B, Richter S and Carlsson J. : Heterogeneity in antigenic expression and radiosensitivity in human colon carcinoma cell lines. In vitro cellular & developmental biology : journal of the Tissue Culture Association 1991, (27A) 12 900–906.

34. Johnson VG, Schlom J, Paterson AJ, Bennett J, Magnani JL and Colcher D. : Analysis of a human tumor-associated glycoprotein (TAG-72) identified by monoclonal antibody B72.3. Cancer research 1986, (46) 2 850–857.

35. Zimmermann B, Gesell T, Chen D, Lorenz C and Schroeder R. : Monitoring genomic sequences during SELEX using high-throughput sequencing: neutral SELEX. PloS one 2010, (5) 2 e9169.

36. Thomas SN, Zhu F, Schnaar RL, Alves CS and Konstantopoulos K. : Carcinoembryonic antigen and CD44 variant isoforms cooperate to mediate colon carcinoma cell adhesion to E- and L-selectin in shear flow. The Journal of biological chemistry 2008, (283) 23 15647– 15655.

37. Sheer DG, Schlom J and Cooper HL. : Purification and composition of the human tumor- associated glycoprotein (TAG-72) defined by monoclonal antibodies CC49 and B72.3. Cancer research 1988, (48) 23 6811–6818.

38. Ruhaak LR, Miyamoto S and Lebrilla CB. : Developments in the identification of glycan biomarkers for the detection of cancer. Molecular & cellular proteomics : MCP 2013, (12) 4 846–855.

